# Helical Reconstruction of Amyloids in cryoSPARC

**DOI:** 10.1101/2025.10.03.680389

**Authors:** Jan-Hannes Schäfer, Robert T. O’Neill, Joseph P. Donnelly, Evan T. Powers, Jeffery W. Kelly, Gabriel C. Lander

## Abstract

Amyloid-mediated proteotoxicity underlies more than 50 human diseases. Cryo-electron microscopy (cryo-EM) analyses have yielded hundreds of in vitro and patient-derived amyloid structures, establishing direct links between filament morphologies and specific pathological conditions. Despite the growing popularity of the processing software cryoSPARC for single-particle analyses, RELION remains the dominant software platform for performing helical reconstruction of amyloid structures, highlighting an area for further development. Here, we present comprehensive processing guidelines for helical reconstruction of helical amyloids using cryoSPARC. Through systematic re-processing and validation of publicly deposited datasets, we demonstrate current capabilities and identify key limitations, emphasizing the need for amyloid-specific parameter optimization within cryoSPARC workflows. Our findings showcase a potential for developing unsupervised processing workflows to meet the demanding throughput requirements of time-resolved in vitro studies and largescale compound screening initiatives, thereby accelerating therapeutic drug development. Ultimately, our goal is to shift the focus of amyloid cryo-EM from computationally intensive processing challenges toward addressing fundamental biological questions that enhance our capacity for treatment discovery.

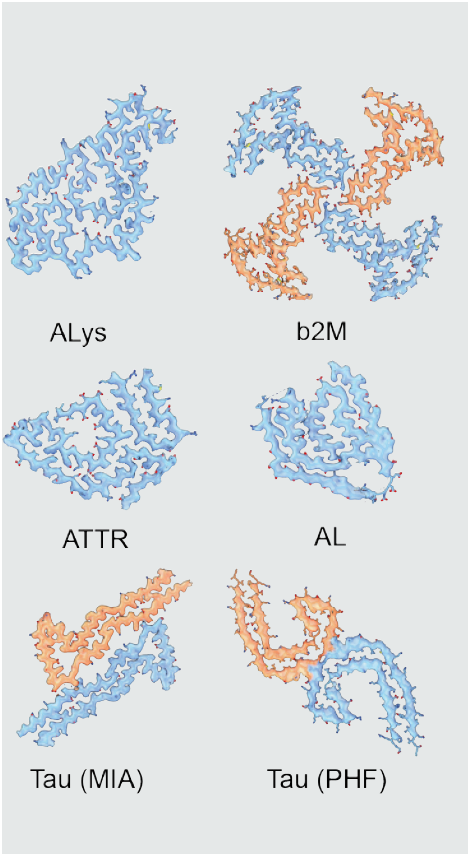

## Introduction

Many macromolecules assemble into filaments with helical symmetry to enable essential cellular functions including motility, morphological organization, and macromolecular trafficking ^1^. While these universal assemblies typically increase protein stability and allow responsiveness to physiological changes through directed assembly and disassembly ^2^, aberrant protein homeostasis can result in pathological fibril formation following irreversible conformational changes. The resulting amyloid assemblies are characterized by stable cross-β-strand arrangements and have a propensity to accumulate into deposits called plaques. Such plaques are implicated in over 50 human diseases, including Alzheimer’s disease (AD), Parkinson’s disease (PD), and transthyretin amyloidosis (ATTR) ^3,4^. Since amyloid fibrils are generally recalcitrant to crystallization, and are too large for NMR analysis, cryoelectron microscopy (cryo-EM) has emerged as the primary method for resolving the molecular details of amyloidogenic disease pathologies. While today’s single-particle workflows can achieve true atomic resolution ^5^, helical reconstruction differs substantially from the traditional workflows used for single particle analyses of macromolecules, requiring specialized processing procedures for accurate structure determination. Helical symmetry optimization tools were previously implemented in RELION to specifically target amyloids, which enabled the first reconstruction of patient-derived tau fibrils ^6,7^. These initial studies led to a flurry of ex-vivo amyloid structures, identifying a link between amyloid folds and underlying disease pathologies and accelerating our understanding of these devastating diseases ^8–10^. Amyloids present unique structural challenges, as they usually present as helical assemblies with constrained screw operators, shallow helical rise values (Δz), and narrow twist ranges (ΔΦ) (Fig.1A). This distinctive symmetry, combined with substantial structural polymorphism observed in ex vivo samples, makes automated filament picking and accurate initial model generation particularly challenging ^11^. Additional complexity arises from substantial structural polymorphism (Fig.1B): Differences in peptide-packing result in polymorphs at the protofilament level, while ultrastructural differences arise from varying numbers of interacting protofilaments within the amyloid fibril. Compositional polymorphs result from ultrastructures of different amyloid proteins and non-amyloid components ^12^. Posttranslational modifications such as glycosylation, disulfide bridges in light-chain amyloids ^13^, or α-synuclein phosphorylation in PD ^14^ add another level of heterogeneity (Fig. 1C). These structural complexities have necessitated amyloid-specific cryo-EM processing tools, which have been successfully developed and implemented within RELION ^7,15–17^. While the Electron Microscopy Data Bank (EMDB) currently reports over 12,700 RELION-generated reconstructions compared to approximately 9,600 from cryoSPARC ^18^, the majority of helical reconstructions — including nearly all amyloid structures — have been processed using RELION (https://www.ebi.ac.uk/emdb/emsearch/charts/, Fig. 1D). Of the 352 amyloid reconstructions documented in the Amyloid Atlas ^19^, only bacterial PSMα1 polymorphs (EMD-43835) ^20^, FapC (EMD-49649) ^21^, and one ex vivo light-chain amyloid reconstruction (EMDB-70557) ^22^ have been processed using cryoSPARC. One possible reason for this discrepancy stems from a user’s greater ability to tightly control for parameter optimization within RELION, whereas cryoSPARC prioritizes generalized processing guidelines. Effective software development relies on community involvement and feedback. To expand cryoSPARC into a truly universal processing pipeline capable of handling both single-particle analysis and helical reconstruction, comprehensive guidelines for reconstructing amyloids need to be established and formalized. This work aims to systematically evaluate the current capabilities of cryoSPARC (v4.7) for resolving amyloid structures while identifying key limitations in existing processing workflows. We use five RELION-processed EMPIAR datasets as case studies and ground truth references to cross-validate reconstructions generated within cryoSPARC. Our comprehensive analysis guides users through the complete processing pipeline: from initial assessment of collected micrographs and rational selection of helical symmetry parameters, through classification of heterogeneous samples and reconstruction of homogeneous polymorphs, to final validation using established community-driven metrics. By providing these systematic guidelines and benchmarks, we hope to accelerate the processing of challenging helical assemblies and unlock new structure-driven insights into amyloid disease pathology and therapeutic intervention strategies.

**Figure 1.**
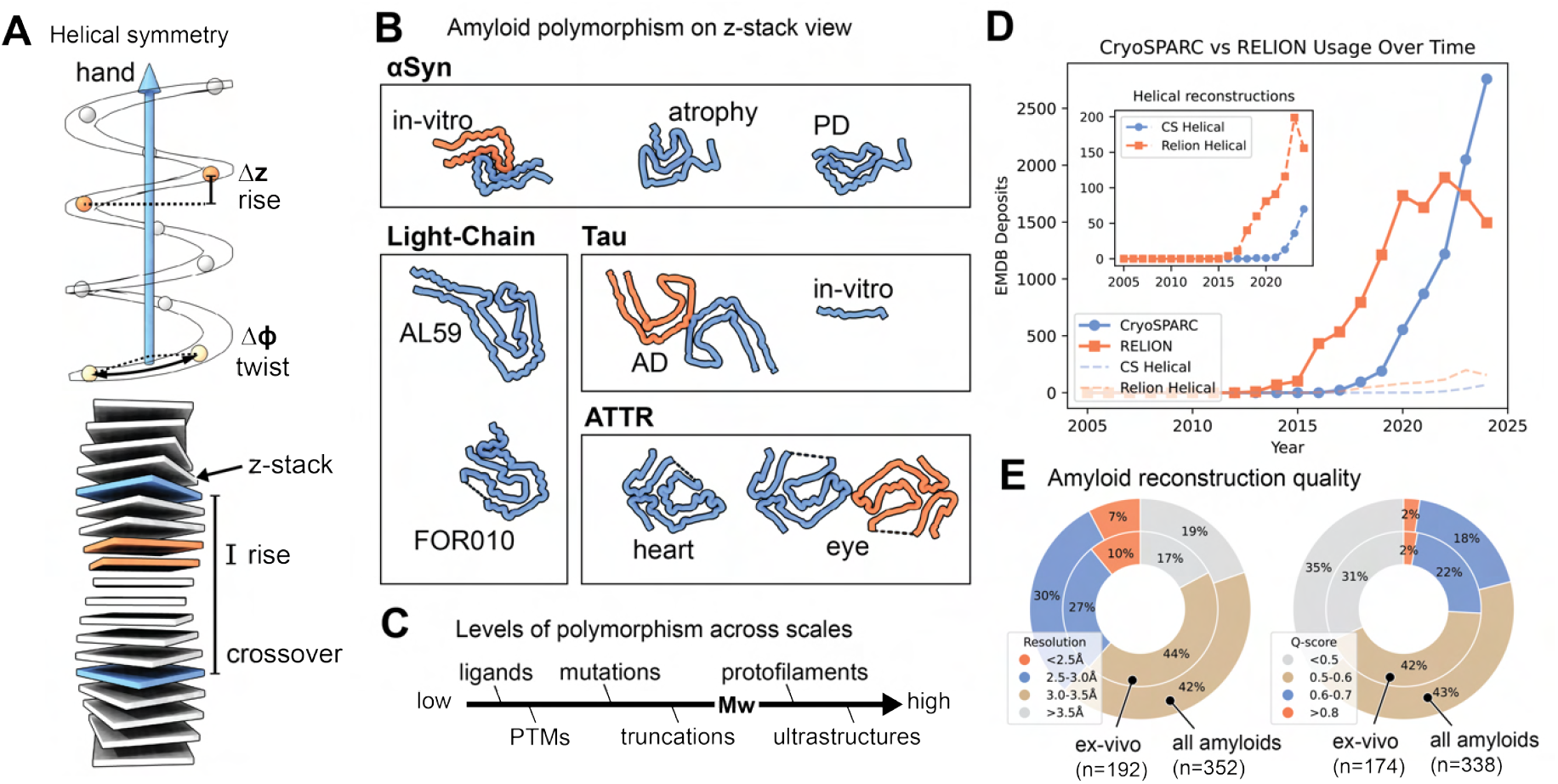
Architecture, diversity, and reconstructions of amyloids. (**A**) Schematic representation of helical symmetry parameters: helical rise (Δz, orange) and helical twist (ΔΦ, tan) with central helical z-axis (blue). Arrow indicates helix handedness. Amyloid z-stack model showing rise (orange) and crossover distance (blue). (**B**) Cross-βpeptide arrangements of amyloid polymorphs grouped by amyloid protein. Single-chain polymorphs shown in blue; double-chain polymorphs in orange/blue. AL59 and FOR010 represent patient identifiers from light-chain amyloidosis cases. (**C**) Contributors to amyloid polymorphism organized by molecular weight (Mw), including post-translational modifications (PTMs). (D) Cumulative EMDB depositions categorized by processing software: RELION (orange) and cryoSPARC (blue). Data for 2025 omitted for annual comparison consistency. Helical reconstructions highlighted in inset. (E) Validation metrics for deposited amyloid reconstructions using Amyloid Atlas data. Q-scores and resolutions at FSC 0.143 for ex vivo and all annotated amyloid entries displayed as nested pie charts.

## Results

### Processing guidelines for amyloids using cryoSPARC

The following approaches for reconstructing amyloids in cryoSPARC were identified through systematic parameter optimization and provide a foundation for target-specific adjustments, though universal applicability may vary. A comprehensive workflow is depicted in Fig. 2.

**Figure 2.**
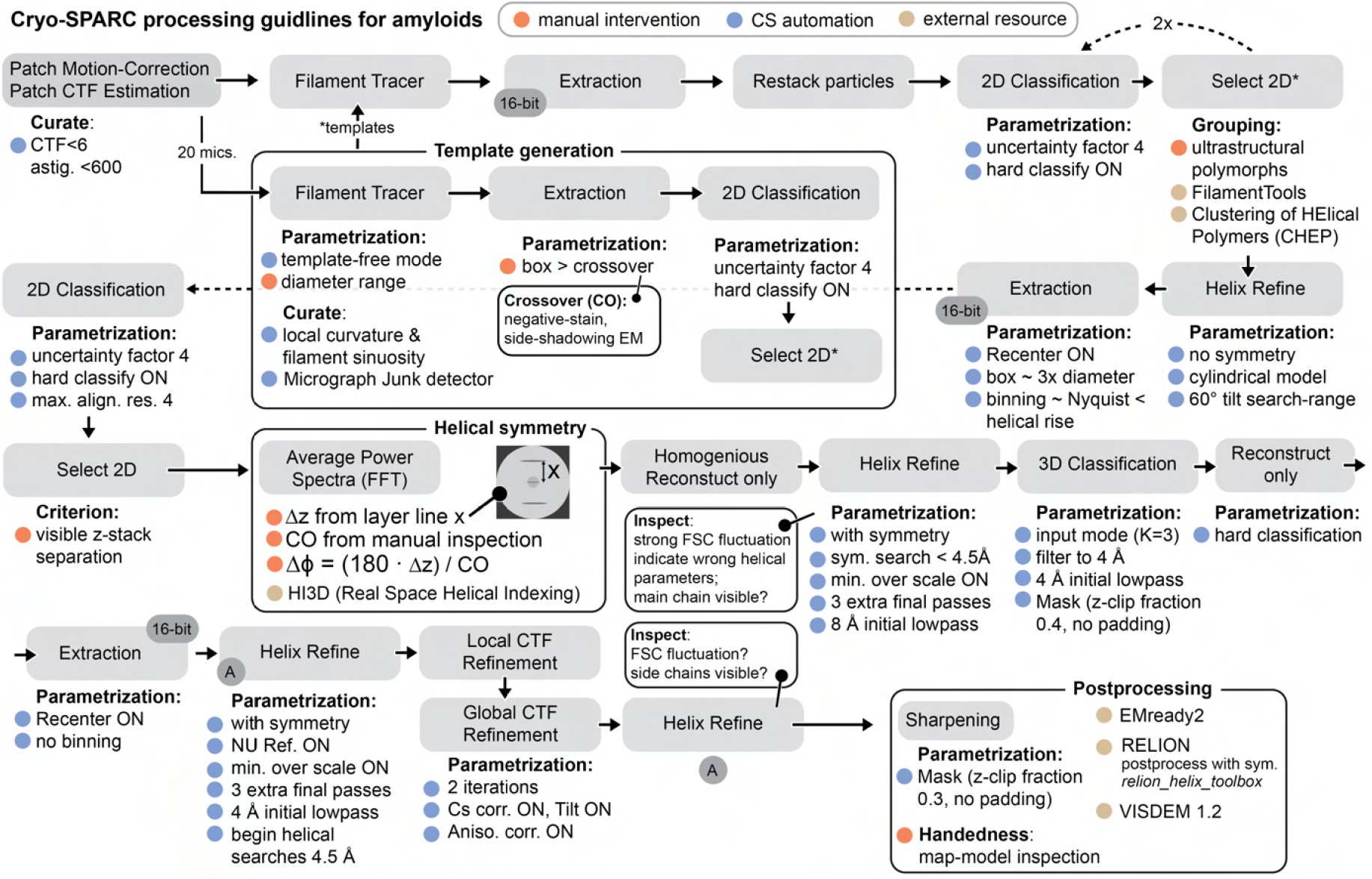
Processing guidelines for amyloids using cryoSPARC. Individual cryoSPARC job types are displayed in grey boxes, with recommended non-default parameters marked by blue bullets. Steps requiring manual intervention or supplementary experiments are denoted by orange bullet points, and external resources are denoted by tan bullet points.

#### Preprocessing and initial considerations

For image pre-processing, beam-induced motion was corrected using Patch Motion Correction, followed by Patch CTF Estimation. Amyloids typically accumulate in thicker ice regions near grid hole edges, necessitating less stringent selection criteria than conventional single-particle analysis. We retained micrographs with CTF fits above 6 Å and discarded those with astigmatism values exceeding 600 Å. Picking templates were generated using template-free Filament Tracing on 20 representative micrographs. The tracer identifies helical filaments as short, straight, partially overlapping segments using an approach adapted from SPRING ^23^. These extracted segments serve as input for iterative helical real-space reconstruction (IHRSR) ^24^. Both RELION and cryoSPARC implement the IHRSR algorithm, segmenting filaments into overlapping short segments that can be averaged while preserving the ability to separate heterogeneous populations. This segmentation approach is feasible because single projections of helical assemblies contain complete information for reconstructing the asymmetric unit ^24^.

#### Filament Tracing to 2D Classification

Filament tracing efficiency improved when diameters slightly below actual filament dimensions were selected, combined with increased standard deviation values for Gaussian blur (0.4-0.5) to enhance filament contour prominence. To separate visually distinct polymorphs, the initial extraction box should exceed the approximate crossover distance, which can be estimated from high-defocus cryo-EM micrographs or negative-stain transmission electron microscopy pre-screening. Two rounds of 2D classification effectively generate well-aligned class averages that enable separation of different filament types. Optimal results were achieved using a large number of classes (K=100) with an increased initial uncertainty factor (e.g., 4) and 40 expectation-maximization iterations, enabling hard classification during the final iteration. This approach effectively reduces per-class heterogeneity, as evidenced by high effective sample size (ESS) values approaching unity and Fourier ring correlation (FRC) values near the Nyquist frequency.

#### Initial Reconstruction Generation and Polymorphism Management

Unlike RELION, cryoSPARC does not support initial model generation with helical priors using projection overlaps across entire helical crossovers ^7^. Instead, cryoSPARC generates cylindrical starting models through its Helical Refinement job. Selecting homogeneous projections that span an entire crossover distance proves most efficient for initial model generation (Fig. S1). Using smaller extraction boxes may produce mixed starting models from conflicting projection sets. Similar to RELION’s expectation-maximization optimizer, the initial volume must fall within the convergence search range to generate accurate final reconstructions ^7^. Separating inherent heterogeneity (polymorphism) requires manual grouping of matching 2D projections in cryoSPARC. Advanced tools such as FilamentTools for bi-hierarchical classification of filament 2D projections in RELION5^16^ or Clustering of Helical Polymers (CHEP) ^25^ are not yet available in cryoSPARC. Consequently, users must manually select class averages representing suspected polymorphs, with inherent risk of false-positive selections. The initial Helical Refinement should proceed without imposing axial or screw symmetry constraints, without limiting shifts along the helical axis, and with maximum out-of-plane tilt searches set to 60°.

#### Symmetry Parameter Determination

While large extraction boxes with high binning enable initial amyloid identification and rough separation of protomer-level polymorphs, helical segments should be re-extracted using smaller box sizes (approximately 300 Å or three times the amyloid diameter) with reduced binning to resolve β-stack separation. This approach typically matches the Nyquist frequency to values below the expected separation distance or helical rise of amyloids (approximately 4.8 Å). To select projections displaying visible stack separation, segments were re-extracted, re-centered, and subjected to 2D classification using maximum alignment resolution of 4 Å, initial class uncertainty factor of 4, and hard classification enabled during the final iteration Fig. S1. In cases of a sharp FSC-falloff near the resolution-range of the rise (4-5 Å), the half-maps may be shifted along the Z-axis. An established route to overcome Z-shifted half-maps is to manually align the half maps (e.g. in ChimeraX) and replace the shifted one in the cryoSPARC job-directory prior to subsequent refinements ^11,17^. The Average Power Spectra job type applied to well-resolved projections enables determination of helical rise from prominent layer lines. By measuring the distance from the meridian to the layer line of interest (d/px), the rise (Δz/Å) can be calculated as: Δz = (box size)/(d) × 2 × (pixel size). Approximating helical twist (ΔΦ) requires the crossover distance, which must be estimated from low-pass filtered micrographs or complementary methods such as negative-stain transmission electron microscopy or atomic force microscopy. The twist can then be calculated using: ΔΦ = (Δz × 180°)/(crossover distance). While reciprocal space measurements provide one approach for deriving screw parameters, real-space helical indexing tools like HI3D offer alternatives by using projections of asymmetric helical reconstructions without requiring manual approximations, accessible through its web-based server ^26^. For improved HI3D accuracy, finer sampling parameters should be employed: angular steps of 0.5° and axial steps of 0.2 Å ^27^. Before proceeding to helical refinement, particle quality should be assessed using the “Homogeneous Reconstruction Only” job with enforced helical rise applied to the selected particle subset.

#### Final amyloid reconstruction through refinements and classifications

To assess the accuracy of the approximated helical parameters, helical refinements should be performed with helical symmetry enabled, 8 Å initial low-pass filtering, per-particle scale minimization enabled, and three additional final passes. Maximum out-of-plane tilt should be set to 60°. Symmetry searches should commence only at 4.5 Å resolution to maintain convergence toward separated cross-β stacks. Strong fluctuations in FSC curves with high discrepancies between tight and corrected FSC curves for auto-masking may indicate incorrect helical symmetry (Fig. S1,S2, see Tau-PHF example). For samples where there is substantial compositional or conformational heterogeneity, 3D classification without alignment may facilitate particle separation into distinct structural classes, to enable symmetry search convergence on subsets to achieve meaningful reconstructions. Reported resolution estimates at this processing stage are unlikely to reflect the actual resolution of the structures, and reconstructions should be critically examined qualitatively for main-chain visibility and stack separation. Classification using input mode with three classes (K=3), filtered to 4 Å with a low-pass filter at 4 Å and focused masking (z-clip fraction of 0.4, no extra padding) may successfully separate low-quality particles or distinct polymorphs. A “Reconstruction Only” job with hard classification enabled and enforced rise values should be performed to inspect reconstructions and select consensus structures for further processing. The selected particle stack should then be re-extracted and re-centered without binning, followed by another round of helical refinement with symmetry searches enabled (starting at 4.5 Å), 4 Å low-pass filtering, non-uniform refinement enabled for three additional final passes, and per-particle scale minimization enabled. Local and global CTF refinement should be attempted, as these jobs may further enhance reconstruction quality. However, after each CTF refinement step, subsequent helical refinements should be performed to evaluate improvements. The resulting reconstructions require careful manual inspection, comparing observed density quality against prior reconstructions, rather than relying on the reported FSC-based resolution estimates, to ensure meaningful structural interpretation. Global CTF refinement with three iterations should include both tilt and spherical aberration correction. In our testing, reference-based motion correction was not performed since not all selected datasets included movie data, though it may provide further resolution improvements where movie frames are available. Finally, reconstructions can be post-processed using B-factor sharpening within cryoSPARC, employing masks with z-clip fractions of 0.3 and no additional padding. Alternatively, EMready2^28^ or VISDEM1.2 (https://github.com/gschroe/visdem) can be used to improve reconstruction interpretability and facilitate model building.

Most amyloids are deposited to structural databases as lefthanded assemblies (exhibiting negative twist values), but absolute handedness must be experimentally verified rather than inferred. This is critical because cryo-EM projection images are inherently ambiguous with respect to handedness—a righthanded helix viewed from one direction produces the same 2D projection as a left-handed helix viewed from the opposite direction. This fundamental ambiguity means that helical reconstructions can converge to either the correct or incorrect handedness with equally good agreement to the experimental data. At sufficiently high resolution (typically better than 3 Å), backbone traces can directly inform handedness determination through visualization of peptide backbone chirality. However, for reconstructions with lower interpretability, techniques such as platinum side-shadowing transmission electron microscopy or atomic force microscopy could be used to discern the handedness of fibril ^29^. Even by integrating these approaches, there will remain a degree of ambiguity regarding handedness, and this caveat should be explicitly stated with the description of the structure. Comparison of helical parameters and structural features with database entries, such as those in the Amyloid Atlas ^19^, can also facilitate both model building and handedness verification.

### Reconstructing amyloids to high resolution using cryoSPARC

To establish robust processing guidelines for reconstructing amyloids in cryoSPARC, six EMPIAR datasets were selected for systematic evaluation. These include five RELION-processed depositions and one recent cryoSPARC-processed deposition: ex vivo paired-helical filament (PHF-type) tau, recombinant intermediate-amyloid tau (MIA-type), cardiac transthyretin amyloids (ATTR), recombinant β2-microglobulin variant V27M, ex vivo lysozyme D87G, and ex vivo light-chain Lambda6 amyloids (Table 1). The datasets were selected to represent the diversity of amyloid diseases, origins (in vitro vs. in vivo), helical symmetries, data quality, availability, and dataset sizes.

**Table 1.** EMPIAR datasets of helical filaments. Helical crossover (CO), resolution (*d*). * denotes our own work, processed in cryoSPARC (v. 4.6–7). Right handedness denoted with positive twist values, left handedness with negative twist values. Helical crossover was calculated as 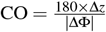.

| EMPIAR | EMDB | $d$ (Å) | PDB | Protein (source) | mic. # | Å/px | Dose (e/Å<sup>2</sup>) | $\Delta z$ (Å) | $\Delta \Phi$ (°) | sym | CO (Å) |
| --- | --- | --- | --- | --- | --- | --- | --- | --- | --- | --- | --- |
| 10940 | 14046 | 2.1 | 7QKL | Tau (MIA) | 331 | 0.824 | 40 | 2.38 | 179.4 | C1 | 778 |
| 10230 | 0259 | 3.2 | 6HRE | Tau (PHF) | 507 | 1.15 | 60 | 2.37 | 179.5 | C1 | 776 |
| 11785 | 18883 | 2.8 | 8R4A | Lysozyme | 2,013 | 1.04 | 49 | 4.78 | -1.4 | C1 | 614 |
| 11383 | 15224 | 2.8 | 8A7Q | b2m-V27M | 611 | 0.94 | 43 | 4.85 | -0.73 | C2 | 1,196 |
| 12815 | 70557 | 2.9 | 9OKA | AL | 7,076 | 0.90 | 40 | 4.86 | 1.2 | C1 | 729 |
| 12217 | 15361 | 2.8 | 8ADE | ATTR | 1,943 | 1.04 | 45 | 4.80 | -1.2 | C1 | 720 |
| 12909* | 71953 | 3.2 | 9PX6 | ATTR wt1 | 1,910 | 0.94 | 50 | 4.85 | -1.26 | C1 | 693 |
| 12911* | 71960 | 3.4 | 9PX7 | ATTR wt2 | 2,426 | 0.94 | 50 | 4.85 | -1.28 | C1 | 682 |
| 12912* | 71962 | 3.0 | 9PX9 | ATTR V122I | 2,326 | 0.94 | 50 | 4.85 | -1.24 | C1 | 704 |

Following the processing guidelines described above, light-chain, transthyretin, lysozyme, tau (MIA) and β2-microglobulin amyloids were successfully reconstructed with high confidence, showing well-resolved side chains (Fig. 3A, Fig. S1S2). However, the PHF tau dataset failed to yield a correct reconstruction (Fig. 3A), highlighting the common problem of convergence to incorrect local refinement minima ^7^. While β-stack separation for ex vivo PHF tau was clearly observable in 2D class averages (Fig. S1), refinement did not converge on correct helical parameters, resulting in an incorrect reconstruction exhibiting signs of overfitting (Fig. S1). Despite extensive iterative refinement attempts with various starting models and symmetry parameters, we were unable to obtain a reliable reconstruction of the PHF tau structure using cryoSPARC. This limitation likely reflects the challenges inherent to complex polymorphic structures rather than fundamental inadequacies of the platform, suggesting that further methodological development could enable successful reconstruction of such challenging targets. In work that is currently under revision, we characterized cardiac ATTR filaments from three different patients (wild-type and familial V122I variant transthyretin), linking amyloid deposits to microangiopathy and capillary occlusion. These amyloids share the common spearhead fold characteristic of transthyretin amyloids, which we reconstructed to high resolution using the presented workflow (Fig. 3B, Table 1). Thus, our processing guidelines enabled meaningful reconstruction of eight out of nine selected datasets in cryoSPARC. Notably, many amyloids exhibit structural polymorphism ^30^. Based on visible protofilaments in 2D class averages, no oligomer-level polymorphism was observed for β2-microglobulin, lysozyme, ATTR, or tau (Fig. S1). Consistent with the original work, the processed light-chain amyloids assemble into single- and double-protomeric arrangements ^22^, but only single-chain amyloids yielded successful reconstructions. Re-processing identified a pseudo-C2 symmetric protomer arrangement in light-chain amyloids (Figure S1) but did not produce a high-quality reconstruction. This polymorphic arrangement was previously reported for a different Lambda6 germline light chain, which also failed to yield reconstructions suitable for atomic model building. The authors suggested a unique 2-fold rotational symmetry perpendicular to the fibril axis ^31^.

**Figure 3.**
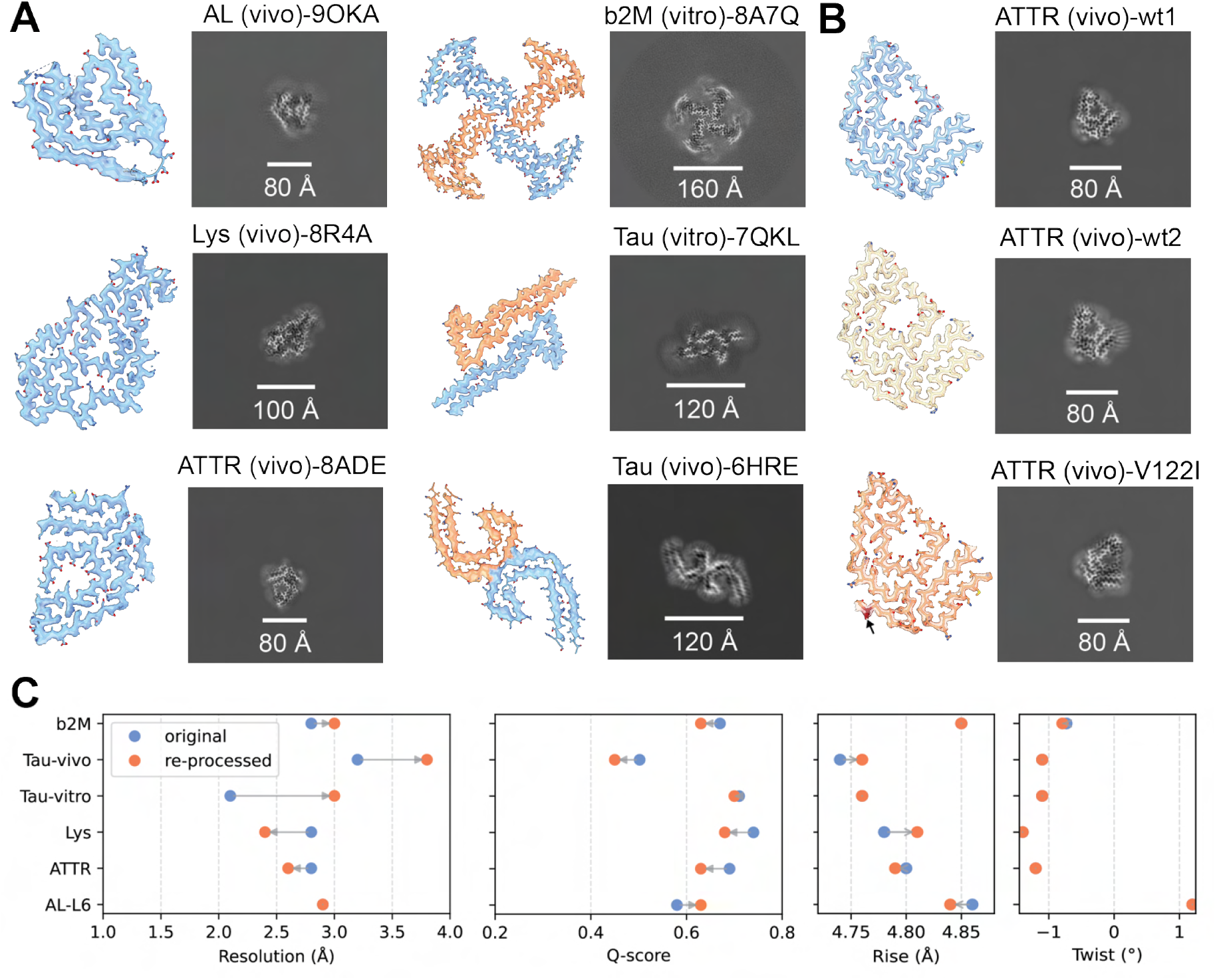
Performance of amyloid processing using cryoSPARC. (**A**) Map-model fits and xy-slices with scale bars for refined reconstructions of amyloid datasets from EMPIAR. Light-chain amyloids (AL, PDB-9OKA), β2-microglobulin variant V27M (β2M, PDB-8A7Q), lysozyme variant D87G (Lys, PDB-8R4A), ex vivo PHF-type tau (PDB-6HRE), transthyretin amyloids (ATTR, PDB-8ADE), and recombinant tau MIA-type amyloids (PDB-7QKL). Single-protofilament polymorphs shown in blue; multi-protofilament polymorphs in blue and orange. (**B**) Cardiac ATTR reconstructions from our previous work showing two wild-type transthyretin samples and familial variant V122I (indicated by arrow). (**C**) Validation of re-processed EMPIAR datasets (orange) compared to original depositions (blue, from Table 1) with resolutions estimated according to an FSC 0.143 cutoff, Q-scores calculated with ChimeraX, and helical symmetry parameters.

### Validating helical reconstructions using cryoSPARC

Helical reconstructions are inherently susceptible to artifacts and subsequent misinterpretation due to the imposition of high-order helical and axial symmetry constraints that inflate the reported resolutions. A recent preprint reported errors in approximately 14% of helical reconstructions deposited in the EMDB, identified through real-space helical indexing and manual data curation ^27^. Given the prevalence of such biases, FSC-based resolution estimation alone cannot reliably assess helical reconstruction quality (Fig. 3C, Fig. S1) and should not serve as the sole validation criterion. Therefore, helical symmetry correctness must be supported by comprehensive map-model assessment, including clear cross-β stack separation, continuous backbone traces, side chain resolvability, and proper stereochemistry ^7^. To quantify the interpretability of our amyloid reconstructions, Q-scores ^32^ were calculated using ChimeraX and compared to original deposited data using corresponding identifiers from Table 1. Q-scores were calculated for single cross-β stacks and plotted in Fig. 3C (detailed values in Table S1). While most reconstructed amyloids support the fitted atomic models with Q-scores above 0.6, the original RELION reconstructions yielded higher interpretability values. The re-processed light-chain AL-L6 from cryoSPARC showed modest improvement compared to the original deposition. Comparison of applied helical symmetry parameters reveals minor discrepancies in helical rise values, potentially contributing to reduced map interpretability. Other established signs of convergence of the refinement to an inaccurate local minimum ^11,17^ are discontinuous density of the backbone trace, negative density between cross β-strands and non-protein fluctuations in local density intensity, which we observed for the PHF tau polymorph (Fig. 3A, Fig. S2).

### Identification of a potential ALys Amyloidosis intermediate

During re-processing of ex vivo lysozyme variant D87G, associated with hereditary systemic lysozyme amyloidosis (ALys) ^33^, non-amyloidogenic fibrils were identified through 2D classification (Fig. 4A). Approximately one-third of micrographs contained these distinct fibrillar structures. Helical refinement using parameters of 37.23 Å rise and 14.17° twist (Fig. S3) yielded a 4 Å resolution reconstruction revealing 10 nm diameter tubes with C6-symmetric protofilament arrangement and a 3.5 nm inner diameter (Fig. 4B). Postprocessing with EMready2 enhanced interpretability, enabling automated model building with ModelAngelo ^34^. Neither human lysozyme C nor annotated human reference proteomes as input produced convincing side-chain placements, necessitating construction of a poly-alanine model for visualization of this unidentified complex (Fig. 4C). The resulting model suggests intra-protofilament packing driven by an elongated β-hairpin motif and inter-protofilament packing through extensive β-complementation lining the tubular pore. While residue-level interpretation remains limited by current resolution, this structure warrants further investigation as it may represent a novel disease-relevant intermediate in ALys amyloidosis pathogenesis or common amyloid-binders like Serumamyloid protein (SAP).

**Figure 4.**
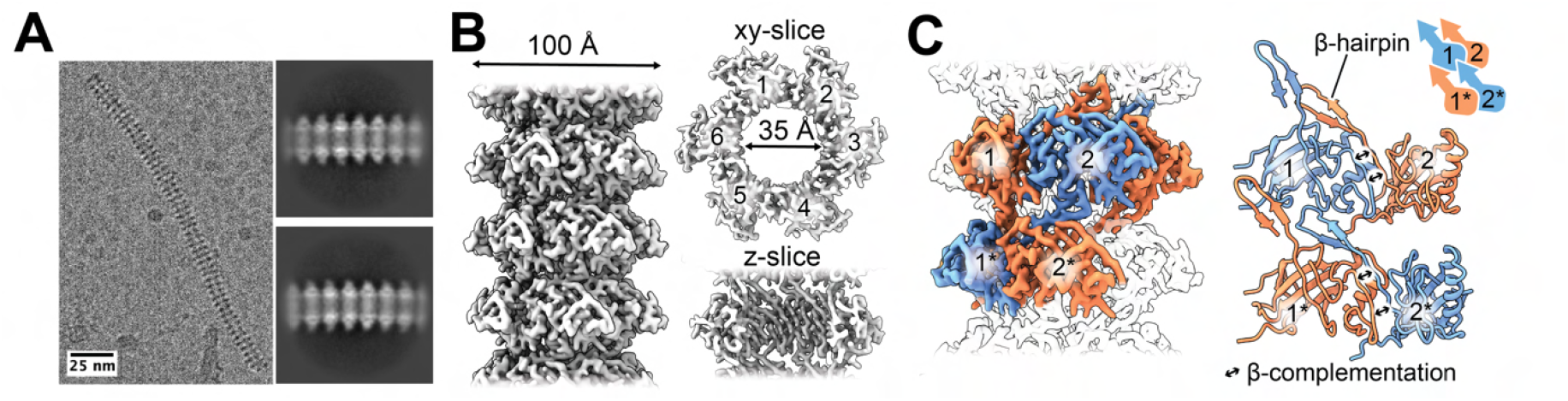
Reconstruction of an unidentified tubular assembly in ex vivo lysozyme amyloids. (**A**) Micrograph from EMPIAR-11785 showing 2D class averages of a non-amyloidogenic tubular fibril (scale bar: 25 nm). (**B**) 4 Å helical reconstruction of 10 nm C6-symmetric tubes with 3.5 nm inner diameter, displayed along the helical axis and as xy- and z-slices. Numerals indicate individual protofilaments. (**C**) Cryo-EM reconstruction color-coded by subunit packing using a poly-alanine model automatically fitted into the density using ModelAngelo. Inter-protofilament contacts through β-complementation are indicated by black double arrows.

## Material & Methods

### Data selection and curation

For database searches, the EMDB REST API (https://www.ebi.ac.uk/emdb/api/) was utilized to automatically extract the Q-score and resolution values for matching EMDB entries of amyloids reported by the Amyloid Atlas ^19^. For testing and validating our processing guidelines in cryoSPARC, we re-processed the amyloid reconstructions from six EMPIAR datasets, listed in Table 1.

### Amyloid processing

All datasets were processed in cryoSPARC v.4.6-7 on NVIDIA Tesla L4 GPUs and Intel Xeon CPU E5-2698 v4 with up to 4 GPUs for multi-GPU CS job-types. Processing of EMPIAR datasets was performed following the established guidelines shown in Fig. 2 and described in the Results section. A comprehensive validation of the individual datasets is shown in Fig. S1,S2 and Table S1. Results were visualized using ChimeraX ^35^ and *ggplot* in Python. Pseudo-screw symmetry in tau amyloids (P2_1_) was converted to a C2 point group for ease of visualization by multiplying the rise value by two and subtracting 180 from the twist value.

### Processing of tubular filaments in EMPIAR-11785

EMPIAR-11785 was processed following the established guidelines shown in Fig. 2. From the 2,013 processed micrographs, about 30% contained non-amyloid segments and were selected for further processing. To increase the number of particles, filaments were traced using a diameter of 40-80 Å without templates and 0.5x gaussian blur. Segments were extracted in a 208 pixel box, Fourier-cropped to 100 pixel and subjected to two rounds of 2D classification with an initial class uncertainty factor of 4 and hard classification during the last iteration. 30K selected segments were re-extracted with recentering in a 256 pixel box and no binning. Helix refine was performed without symmetry using a cylindrical model and a 60° tilt search-range. The resulting reconstruction was used as input in Real-space helical indexing in HI3D with default parameters ^26^, suggesting a twist of 13.74° and a rise of 37.93 Å. These values were used in helix refine with non-uniform refinement (NUR) enabled. To test for higher-order symmetry, Phenix’s symmetry search tool phenix.map symmetry ^36^ was used, resulting in the highest correlation with C6 point-group symmetry. After another round of 2D classification, 21K segments were used in helical refinement with C6, a rise of 38 Å and twist of 13.7°. Helical indexing with the new C6 point group and higher-resolution map suggested a rise of 37.19 Å and twist of 14.21°. The final helical refinement with a 20° tilt search-range resulted in a 4 Å reconstruction (FSC 0.143). The map was sharpened automatically within cryoSPARC using a B-factor of –56 Å 2. The full processing workflow and validation is shown in Fig. S3 and Table S1. Resolvability of the map was increased with postprocessing in EMready2^28^ and utilized for automatic model building with Model Angelo ^34^. Human lysozyme C (Uniprot: P61626) and the annotated human reference proteome (UP000005640) were used as input in FASTA or the HMM search procedure, respectively.

## Discussion

In our previous work on ex vivo ATTR amyloids, we demonstrated cryoSPARC’s capability to reconstruct amyloids from complex tissue extracts to high resolution (Fig. 3B). This success motivated the development of systematic processing guidelines for amyloid reconstruction using cryoSPARC as a viable alternative to RELION. Following our ATTR work, we re-processed deposited amyloid datasets using our established workflow, successfully reconstructing five of six selected amyloid datasets to high resolution, and only one dataset failed to achieve suitable resolution for model building, since it converged onto a local but incorrect minimum (Fig. S2). Notably, our new processing workflow, available as import-ready.json files (https://github.com/schaefer-jh/CS-amyloids/), achieved 2.4 Å global resolution for ALys amyloids (Fig. 3A, Fig. S1) in approximately 12 hours using three GPUs for multi-GPU processing of 2,000 movies. By integrating our recently published deep neural network CryoSift (https://cryosift.org) ^37^, which automatically selects suitable 2D class averages through its integrated mode with cryoSPARC-tools, unsupervised high-throughput processing of amyloids with known helical symmetry becomes feasible in cryoSPARC, potentially accelerating future drug development efforts. While many of the presented non-default processing parameters facilitate reconstruction of homogeneous amyloids and protofilament-level polymorphs in cryoSPARC, reliable classification of polymorphs at the peptide level remains challenging. To enhance cryoSPARC’s capabilities as a complementary platform for amyloid reconstruction, several targeted improvements would expand its utility alongside existing tools like RELION. Key improvements include implementing non-cylindrical starting-model generation using projection-level information from overlapping 2D class averages, similar to *relion helix inimodel2d* in RELION ^7^ or *denovo3d* in Helicon ^38^. Coupled with automatic clustering of 2D class averages, analogous to RELION’s FilamentTools ^16^, this would create more homogeneous particle groups, helping overcome the common problem of convergence to local refinement minima by reducing the search range for initial model generation while minimizing user error in class selection ^7,11^. While large-scale polymorphs such as single and double protofilaments of light-chain amyloids (Fig. S1,S2) were readily classified at the projection level, previous studies on light-chain Lambda6^31^ demonstrated that double protofilaments contain additional peptide-level morphologies that require further 3D classification. These polymorphs and structural intermediates likely play crucial roles in amyloid disease progression ^39^ and have been successfully dissected using time-course cryo-EM experiments ^16^. Separation and high-resolution reconstruction of ex vivo amyloid mixtures is essential for therapeutic development; however, cryoSPARC currently lacks 3D classification with helical rise and twist optimization, making separation of unique but similar filament types difficult or impossible. Implementing classification routines with helical symmetry optimization, both with alignment in Heterogeneous Refinement and without alignment in 3D Classification jobs, would enable broader access to amyloid structure determination using cryoSPARC. Re-processing cryo-EM datasets can reveal unexpected findings, such as vault particles discovered in ex vivo tau samples ^40^. This proved true for the deposited lysozyme amyloids associated with systemic ALys amyloidosis as well. Through 2D classification, we identified previously unreported non-amyloid fibrils, which we reconstructed to 4 Å resolution, revealing a hexameric tubular structure with β-sheets stabilizing inter- and intra-protomer interfaces (Fig. 4C). Due to limited complementary data and resolution constraints, incomplete side-chain information prevented identification of the protein forming these tubular structures. We speculate this helical assembly may represent a disease-relevant intermediate in ALys amyloidosis pathogenesis,, or stacks of serum amyloid P component (SAP), a common binding-partner of amyloids, though further investigation is required for confirmation ^41^. Our study presents current capabilities and limitations of cryoSPARC for amyloid reconstruction using optimized non-default parameters. The established guidelines should enable the reconstruction of diverse amyloids; however, further optimization of amyloidspecific software is needed to advance cryoSPARC’s potential as a universal platform for helical reconstructions.

These community-driven guidelines aim to democratize amyloid structure determination, enabling broader participation in amyloidosis research and accelerating the development of therapeutic interventions.

## Acknowledgements

Funding for this research was provided by a NINDS grant NS095892 to GCL and the German Research Council project number 556478029 to JHS. The authors thank Jean-Christophe Ducom and Charles Bowman at Scripps for computational support, and Sjors Scheres and Danielle A. Grotjahn for critical reading of the manuscript and incisive comments. We thank Olga Gursky and Chad W. Hicks for sharing a pre-release of their EMPIAR-12815 data.

## Data & software availability

All re-processed datasets from the Electron Microscopy Public Image Archive (EMPIAR) were deposited in the Electron Microscopy Data Bank (EMDB). The datasets from Table S1 are available using the following EMDB accession numbers with their respective EMPIAR-ID: ATTR wt1 (EMD-71953, EMPIAR-12909), ATTR wt2 (EMD-71960, EMPIAR-12911), ATTR V122I (EMD-71962, EMPIAR-12912), Unknown (EMD-72138, EMPIAR-11785), AL-L6 (EMD-72157, EMPIAR-19992), ATTR (EMD-72158, EMPIAR-12217), b2M V27M (EMD-72159, EMPIAR-11383), Lys D87G (EMD-72161, EMPIAR-11785), Tau MIA (EMD-72164, EMPIAR-10940), Tau PHF (EMD-72174, EMPIAR-10230). An import-ready.json workflow for cryoSPARC (generated in v.4.7.0) is available on https://github.com/schaeferjh/CS-amyloids/ and can be used per the developers’ instructions (https://guide.cryosparc.com/application-guidev4.0+/workflows).

## Supporting Information

**Figure S1.**
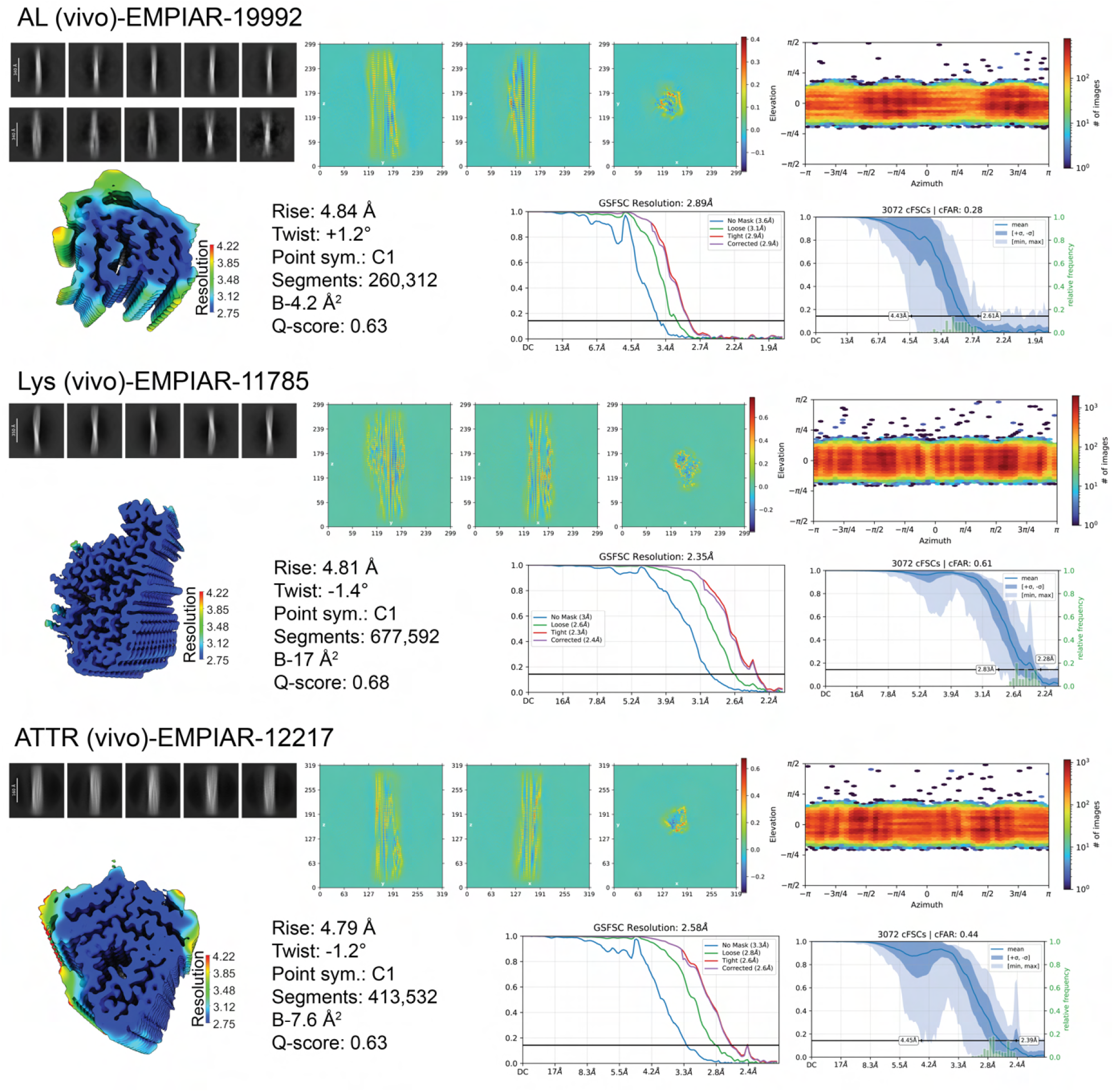
Validation of amyloid reconstructions using cryoSPARC. Comprehensive validation for each EMPIAR dataset includes 2D class averages, real-space slices along major orthogonal planes (zy, zx, xy), viewing directional distribution, local resolution maps (color-coded consistently across all datasets), resolution estimated using gold-standard FSC (cutoff at 0.143), and cFAR reports.

**Figure S2.**
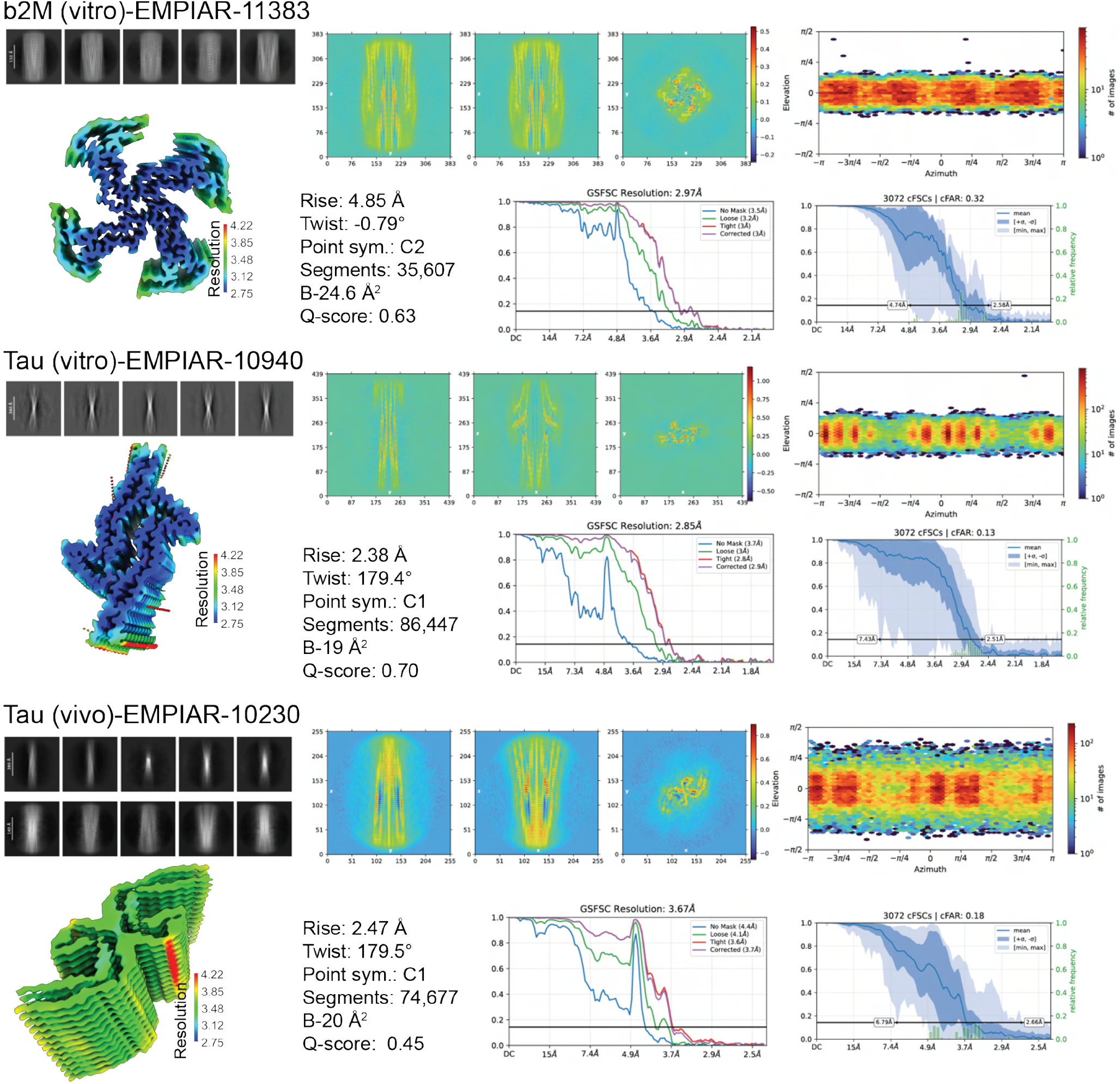

**Figure S3.**
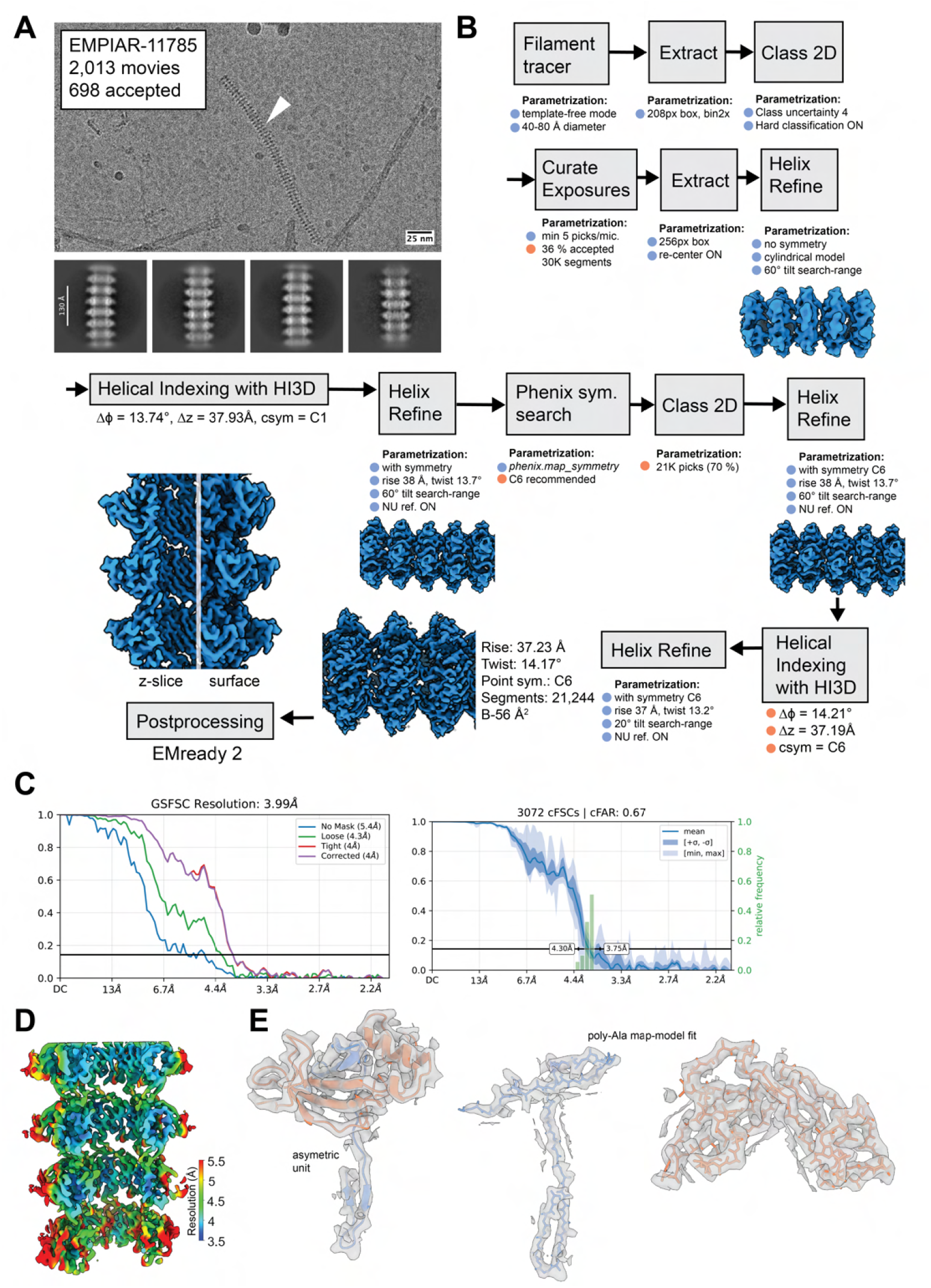
Validation of non-amyloid fibrils. (A) Micrograph and 2D class averages from EMPIAR-11785 positive for the non-amyloid filament (white arrow). 25 nm scale bar on micrograph and 130 Å scale bar on 2D class averages. (B) Processing workflow for helical reconstruction. (C) Gold-standard FSC resolution estimation at 0.143

**Table S1.**
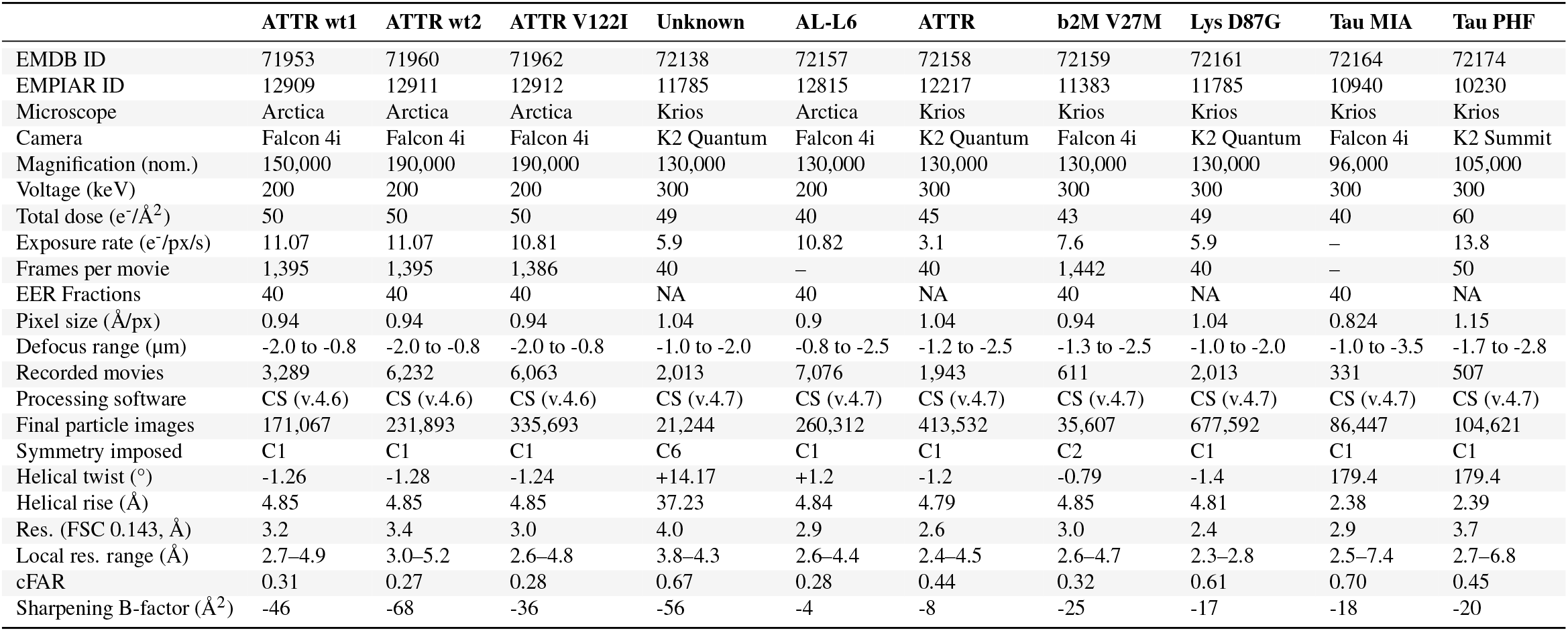
Cryo-EM data collection and image processing of amyloids. (NA, not applicable. -, data not available).

